## Supplementary material for "Intron-loss in Kinetoplastea correlates with a non-functional EJC and loss of NMD factors": Fig. S1

### Figure S1

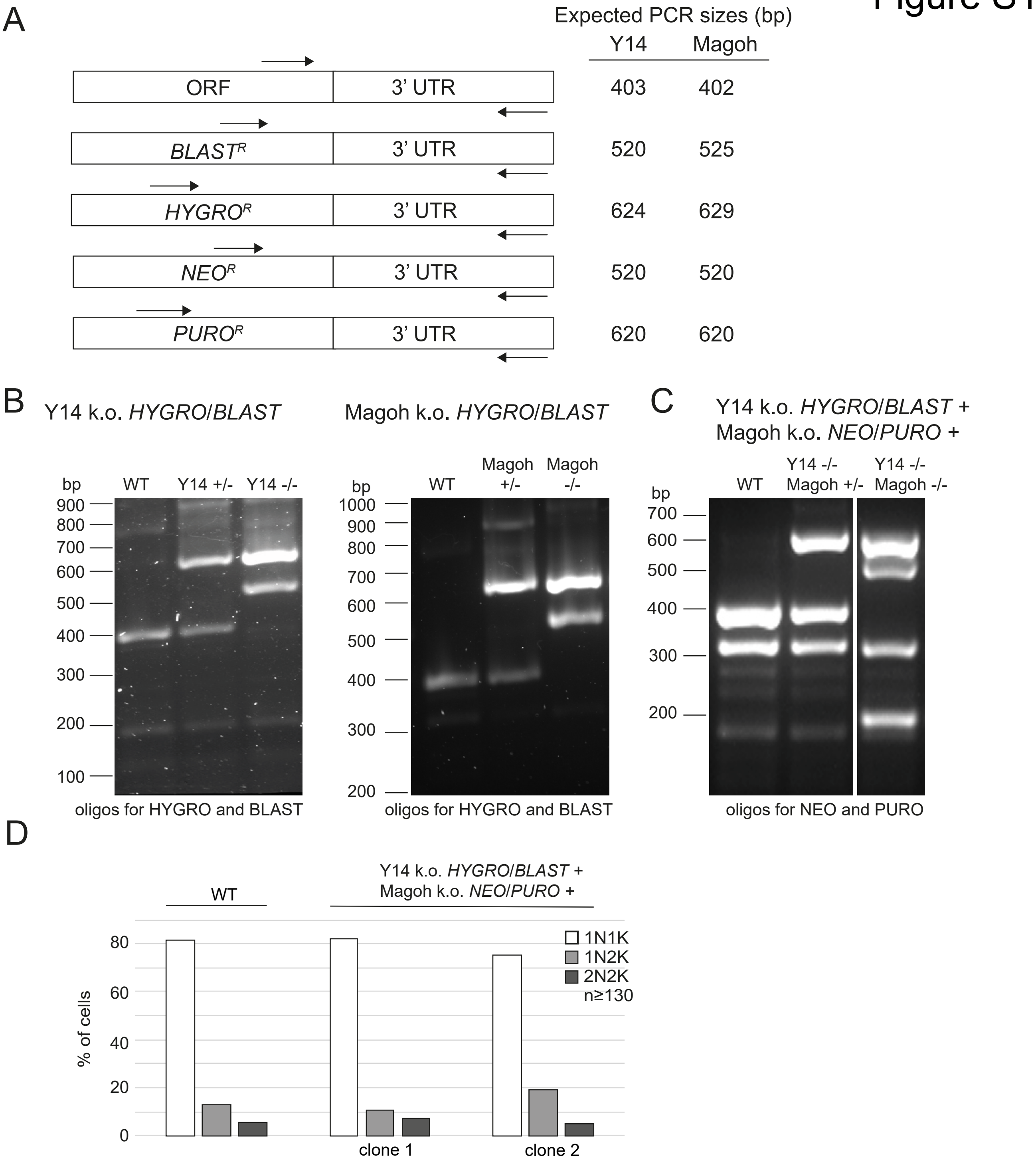


**Figure S1:** Single and double knockout cell lines were generated for *T. brucei* Y14 and Magoh by replacing the endogenous alleles with different resistance markers. For the single knockouts, hygromycin and blasticidine resistances were used for the replacements. The Y14/Magoh double knockout was generated by replacing both Magoh alleles of a Y14 knockout cell line with neomycin and puromycin.

**(A)** Schematics of the wild type allele and the modified alleles, to demonstrate the PCR strategy used for cell line evalution. The expected sizes of the PCR products are indicated for both Magoh and Y14 on the right. **(B)** Individual knockouts were confirmed by PCR, for both Y14 and Magoh. Each PCR reaction contained 4 oligos: forwards oligos specific to the wild type, HYGRO and BLAST and a reverse oligo specific to the 3´UTR of the respective gene. **(C)** The correct replacement of both Magoh alleles by neomycin and puromycin resistance cassettes in Y14 knockout cell was confirmed with a similar PCR strategy. There is an non-specific PCR product of 300 nucleotides. **(D)** Trypanosome cells were stained with the DNA dye DAPI and cells were classified according to their cell cycle stage by counting the number of nuclei (N) and kintoplasts (K) for at least 130 cells. The kinetoplast (the visible DNA of the single mitochondrion) divides prior to the nucleus, allowing a classification into 1K1N, 2K1N and 2K2N cells. No significant differences in the distribution of the different cell cycle stages was observed between wild type cells and the Magoh/Y14 double knockout.

ORF: open reading frame; WT: wild type; k.o.:knockout
