## Supplementary material for "Intron-loss in Kinetoplastea correlates with a non-functional EJC and loss of NMD factors": Fig. S2

### Figure S2

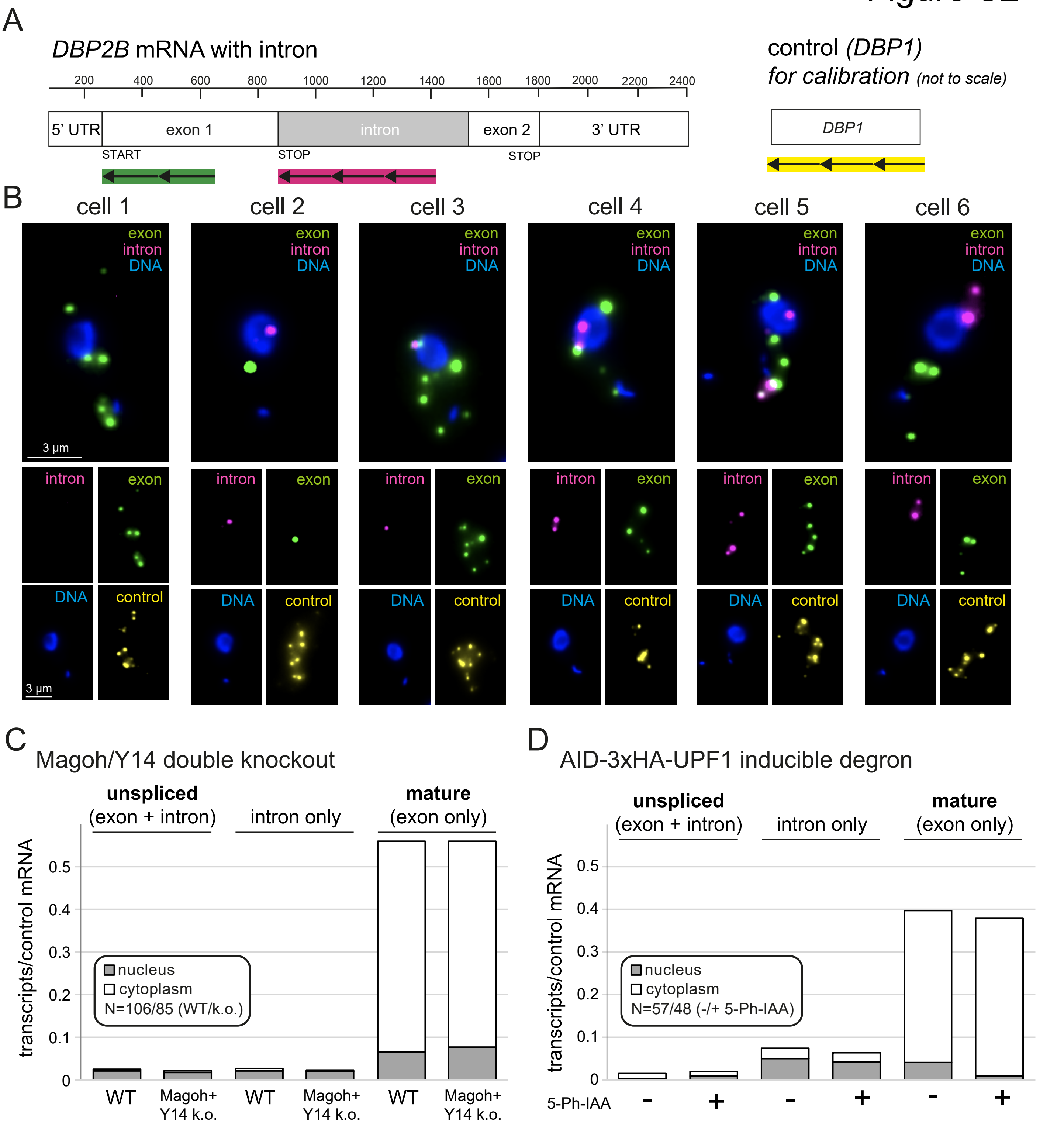


**Figure S2: No significant changes in processing or transport of intron-containing mRNAs in the Magoh/Y14 double knockout and after UPF1 depletion**

**(A)** *DBP2B* is one of two intron containing mRNAs and is here shown schematically prior to intron removal. The positions of the FISH probes used to detect the exon and the intron are indicated.The intronless mRNA DBP1 served as a control for calibration.

**(B)** *DBP2B* exon, *DBP2B* intron and the control mRNA were detected by triple colour single molecule FISH using branched DNA technology of the Affymetrix system as previously described [(1)]. Deconvolved images of a Z-stack (63 slices a 140 nm) are shown as projections (sum-slices) as merged (top) and single channels (bottom), for six representative cells.

**(C and D)** For the Magoh/Y14 double knock-out (C) and the auxin inducible degradation of UPF1 (D) we quantified the amount of unspliced (exon + intron), intron and mature (exon only) *DBP2B* transcripts per control mRNA, in each compartment (nucleus or cytoplasm). For both conditions, we found no significant differences in amount or distribution of the different transcript types in comparison to the control cells (WT or uninduced). In particular, the level of unspliced mRNA did not increase, neither in the nucleus, nor in the cytoplasm.

[(1)] Kramer, S. et al. Parallel monitoring of RNA abundance, localization and compactness with correlative single molecule FISH on LR White embedded samples. Nucleic Acids Res 49, gkaa1142- (2020).
