## Supplementary material for "Intron-loss in Kinetoplastea correlates with a non-functional EJC and loss of NMD factors": Fig. S3

### Figure S3

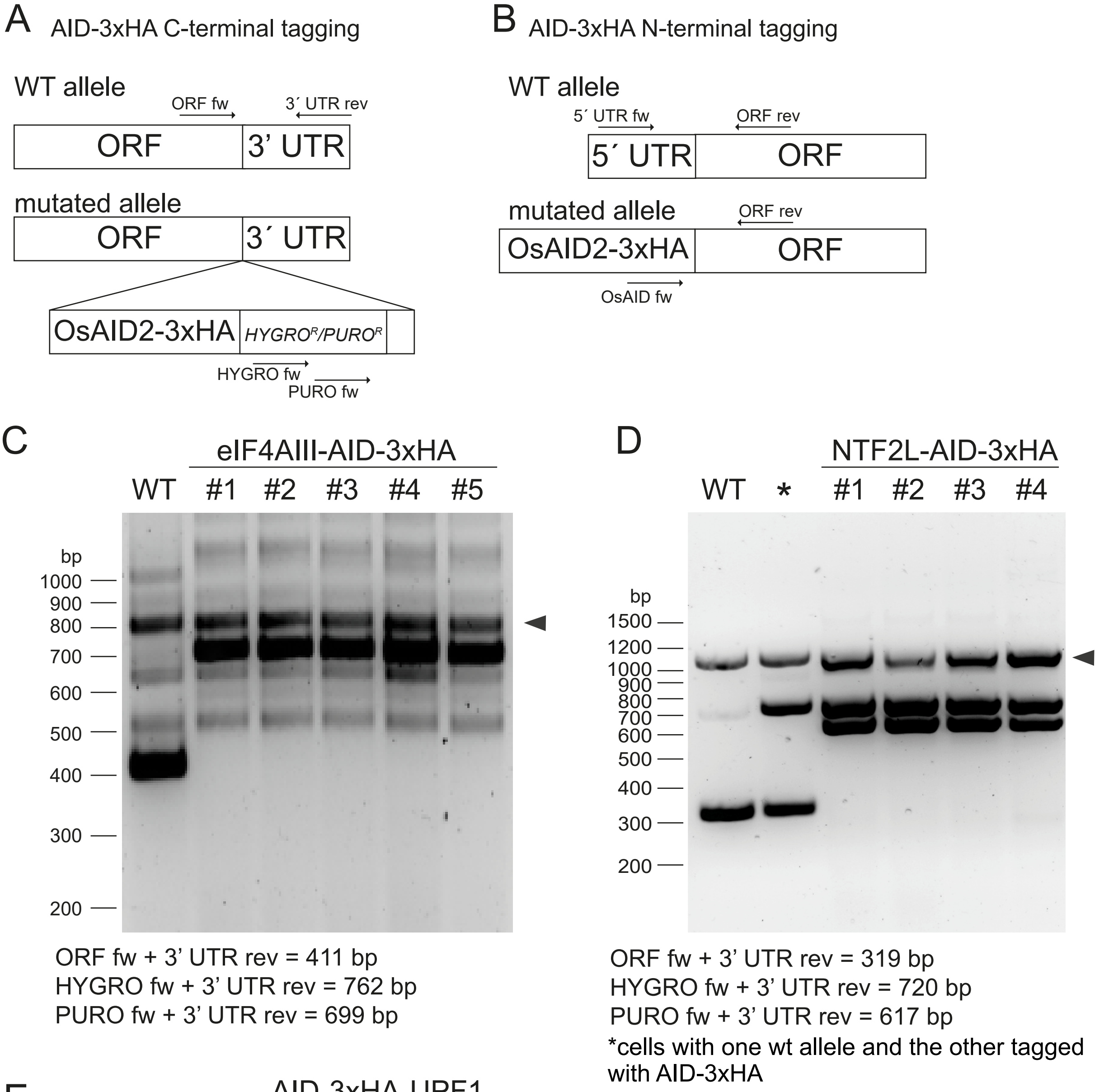


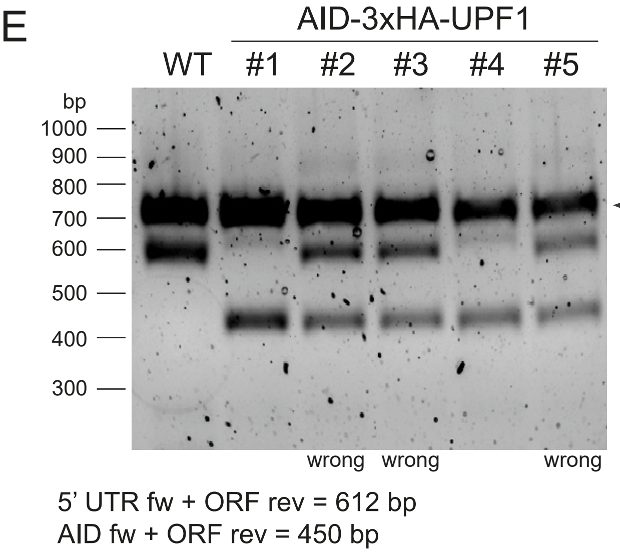


**Figure S3:** The auxin inducible degron system was employed for inducible degration of *T. brucei* eIF4AIII, NTF2L and UPF1. For each gene, both alleles were fused to AID2 either C-terminally (eIF4AIII, NTF2) or N-terminally (UPF1).
**(A-B)** The PCR strategy that was used to evaluate the cell lines and, in particular, to control for the absence of a wild type allele, is schematically pictured for the C- terminal and N-terminal tagging strategy.
**(C-E)** PCR products, using the oligo mixtures indicated below the gels, are shown for wild type cells, several clones (#) of the final cell lines (with both alleles replaced), and for the hemizygote cell line (NTF2L only, marked with *). The expected sizes of the PCR prodcuts are indicated below the gels. Non-specific bands are marked with an triangle. For UPF1, only 2 of the 5 clones tested were correct.
