## Supplementary material for "Intron-loss in Kinetoplastea correlates with a non-functional EJC and loss of NMD factors": Fig. S4

### Figure S4

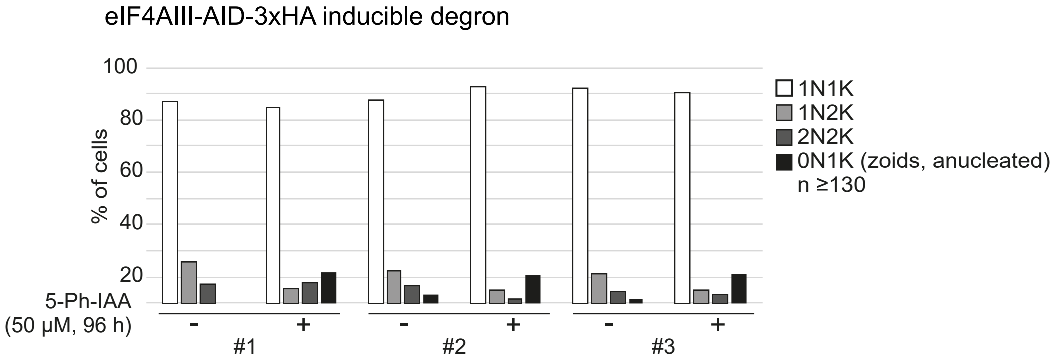


**Figure S4: eIF4AIII depletion causes no specific cell cycle block**
Trypanosome cells were stained with the DNA dye DAPI and cells were classified according to their cell cycle stage by counting the number of nuclei (N) and kintoplasts (K) for at least 130 cells. The kinetoplast (the visible DNA of the single mitochondrion) divides prior to the nucleus, allowing a classification into 1K1N, 2K1N and 2K2N cells. No significant differences in the distribution of the different cell cycle stages was observed upon induction of eIF4AIII depletion, apart from a small increase in anucleated cells.
