## Supplementary figures and images for "Intron-loss in Kinetoplastea correlates with a non-functional EJC and loss of NMD factors"

### Fig. S5

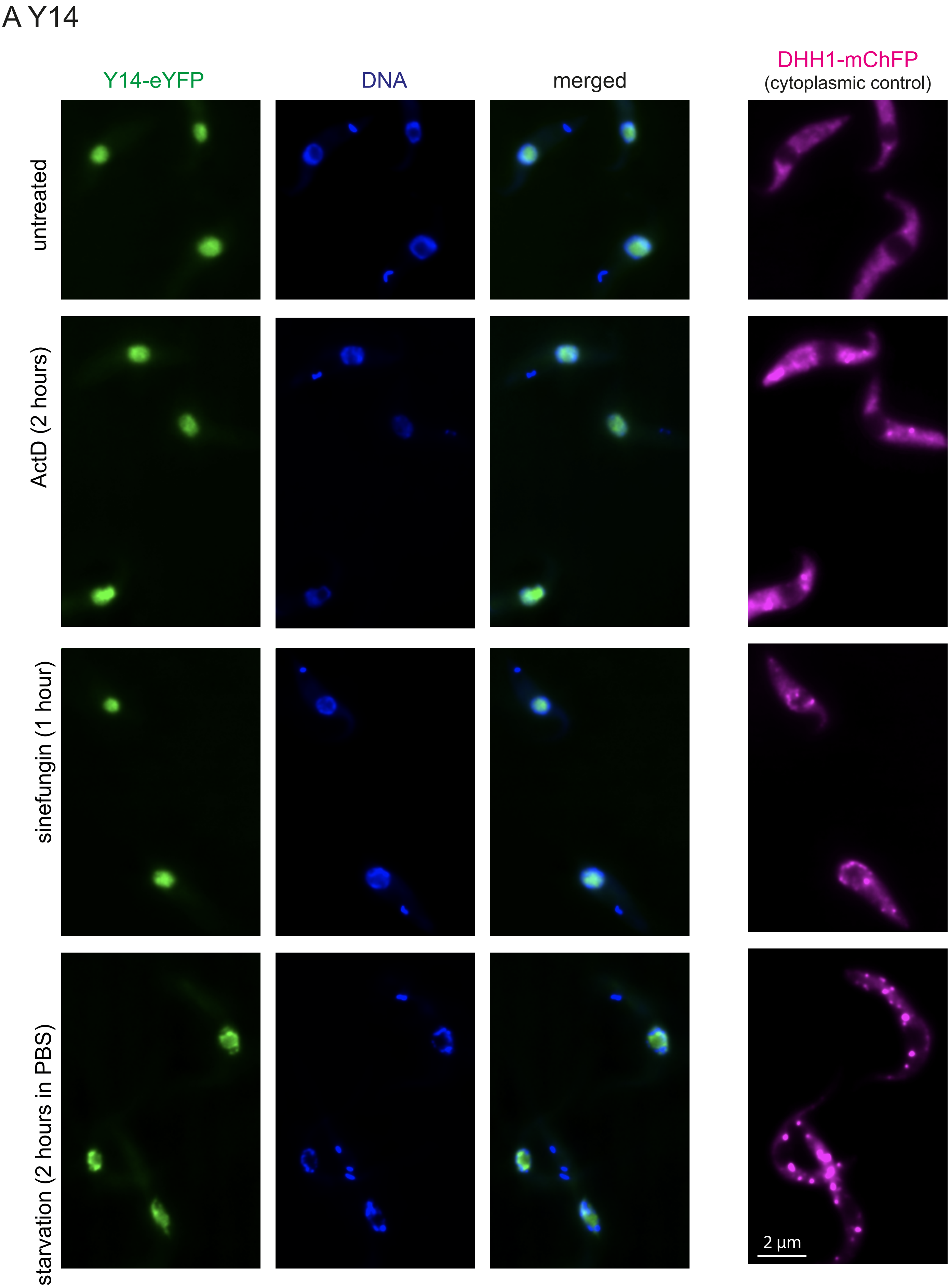

### Fig. S5

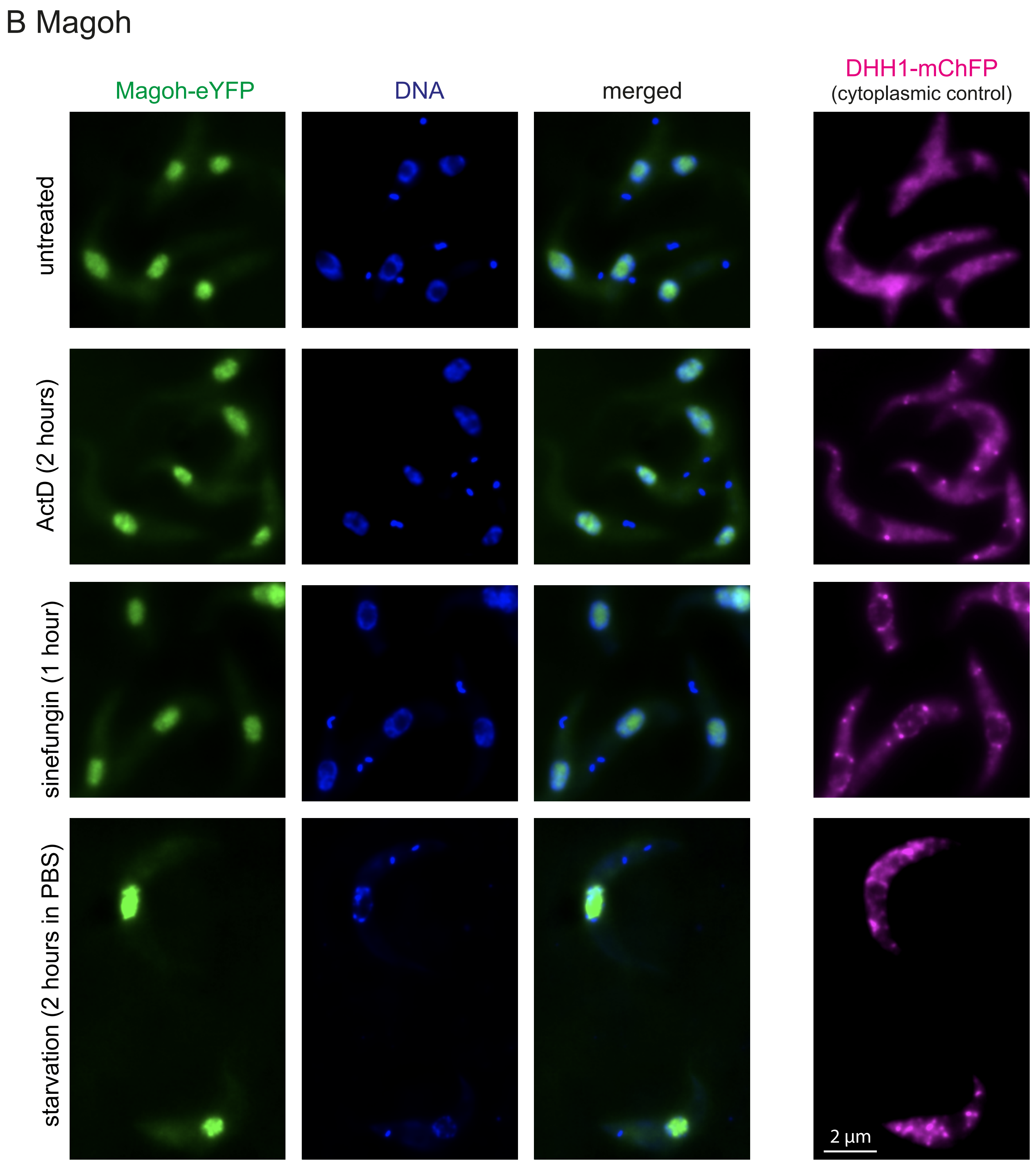
