## Supplementary material for "Intron-loss in Kinetoplastea correlates with a non-functional EJC and loss of NMD factors": Fig. S5

Figure S5

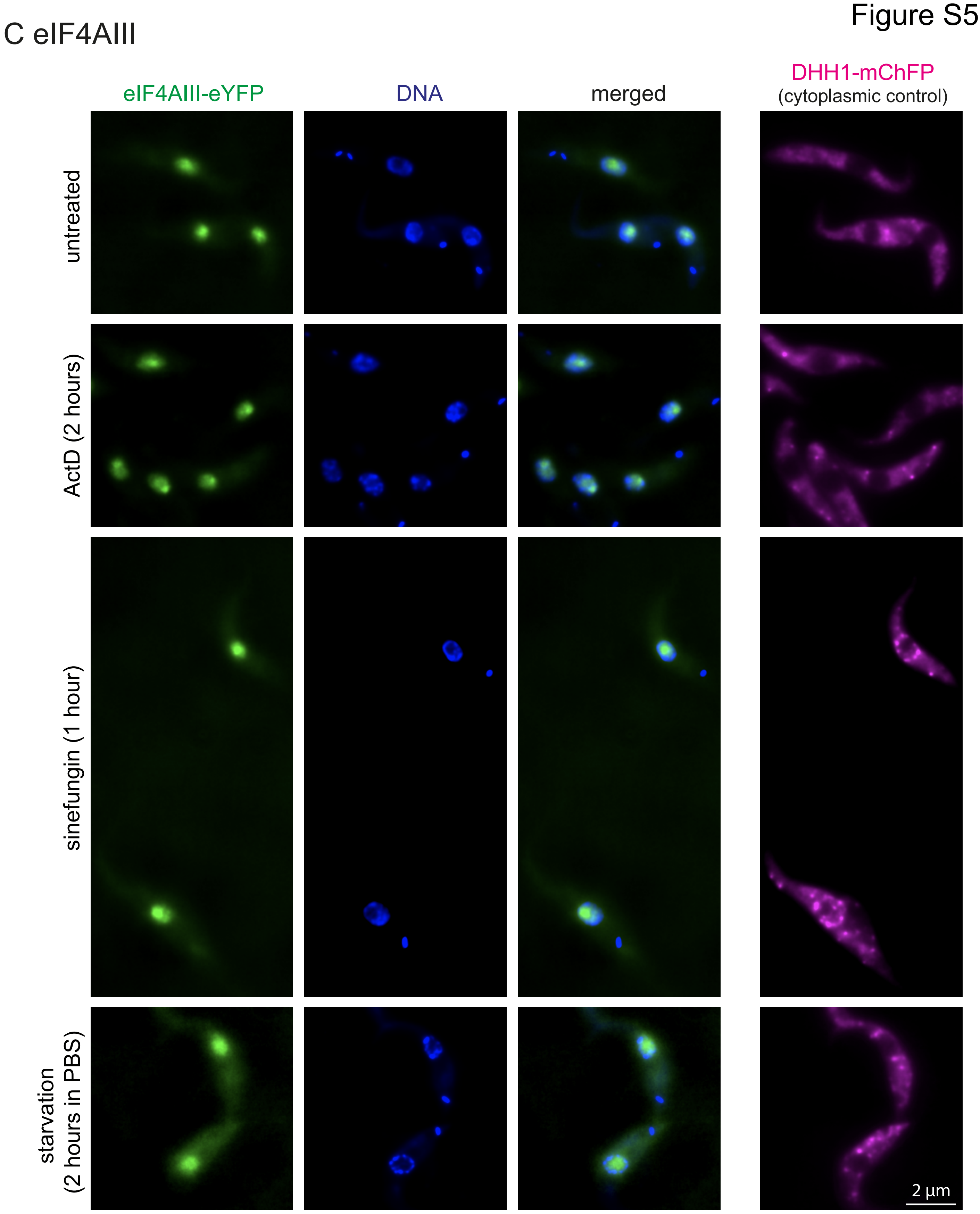


Figure S5: Cells expressing Y14 (A), Magoh (B) or eIF4AIII (C) fused to eYFP in a cell line also expressing DHH1-mChFP were treated as indicated. Projections (sum slices) of deconvolved Z-stacks (75 slices a 140 nm) are shown.
