## Supplementary material for "Intron-loss in Kinetoplastea correlates with a non-functional EJC and loss of NMD factors": Fig.S6

### Figure S6

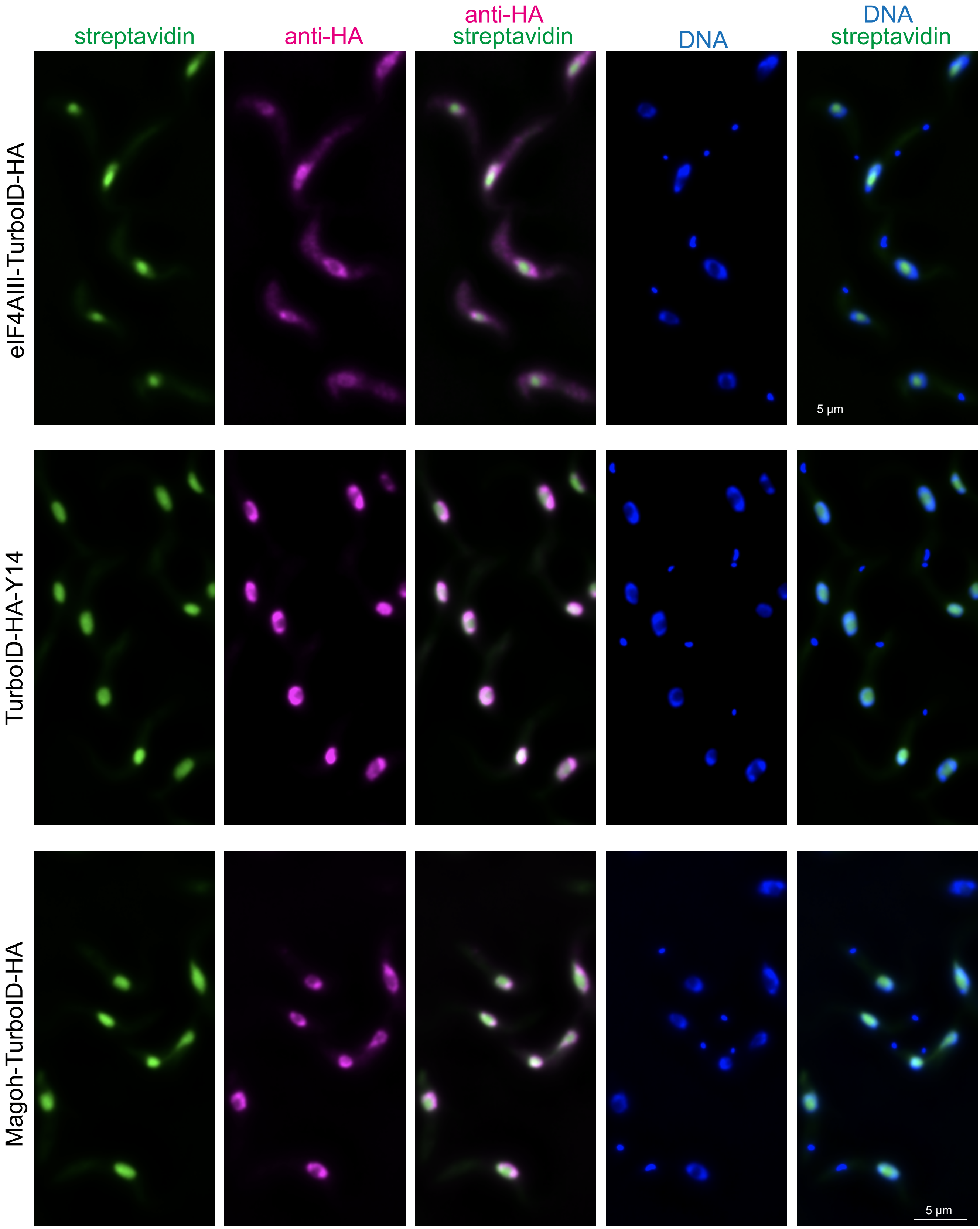


**Figure S6:** eIF4AIII, Y14 or Magoh were expressed fused to TurboID-HA and imaged with anti-HA (shown in pink) and streptavidin-Alexa488. Z-stack projections (sum slices) of a deconvolved Z-stack image (100 slices a 100 nm) are shown.
