## Supplementary material for "Intron-loss in Kinetoplastea correlates with a non-functional EJC and loss of NMD factors": Fig. S7

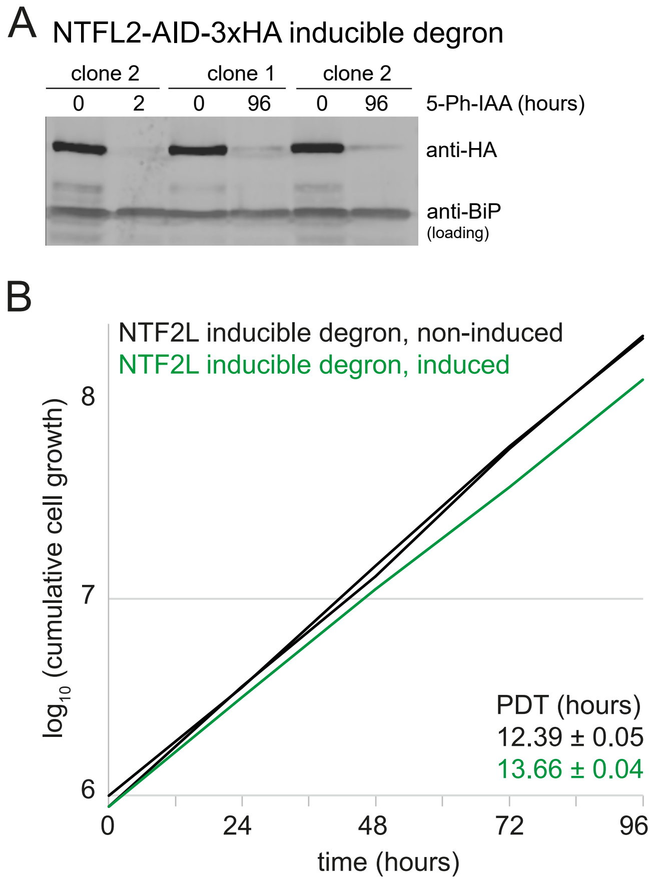


**Figure S7:** Two clonal cell lines (clone 1 and clone 2) were generated for the **depletion of NTFL2 via the AID2 system.**
**A)** Western blot loaded with cell extracts of cells incubated with 50 µM 5-Ph-IAA for 0, 2 or 96 hours. NTFL2-AID-3HA protein is detected with anti-HA. Anti-BIP served as loading control. The NTFL2 protein becomes undetectable within 2 hours of induction.
**B)** Growth was monitored for both clones over 96 hours, in the presence and absence of 5-Ph-IAA. The population doubling times (PDT) are only marginally increased upon induction.
