## Supplementary material for "Intron-loss in Kinetoplastea correlates with a non-functional EJC and loss of NMD factors": Fig. S8

Figure S6

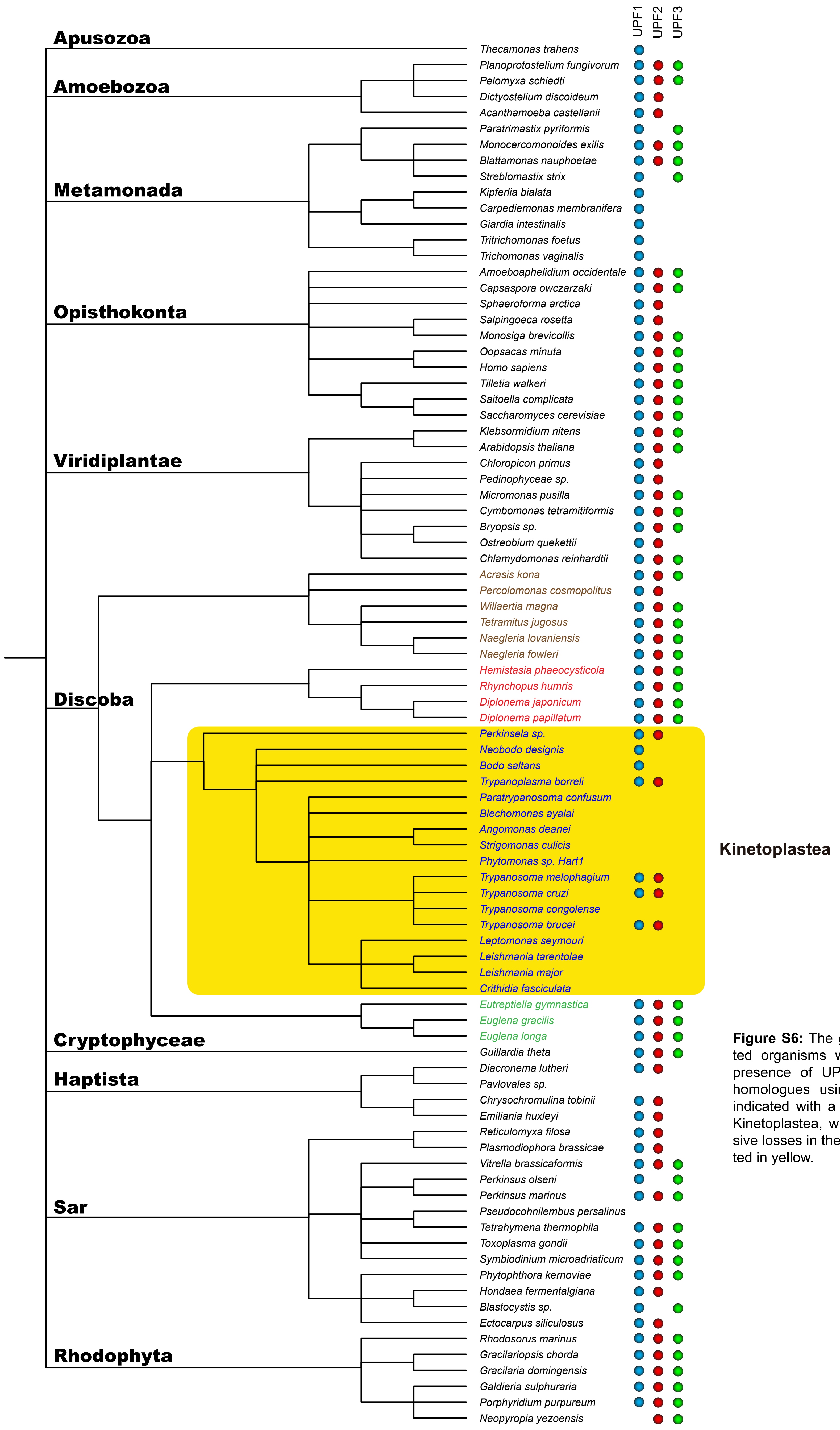


**Figure S6:** The genomes of the indicated organisms were screened for the presence of UPF1, UPF2 and UPF3 homologues using Blast; presence is indicated with a ball. The group of the Kinetoplastea, which experience extensive losses in these proteins, is highlighted in yellow.
